# Nuclear gene transformation in a dinoflagellate

**DOI:** 10.1101/602821

**Authors:** Brittany N. Sprecher, Huan Zhang, Senjie Lin

**Affiliations:** Department of Marine Sciences, University of Connecticut, 1080 Shennecossett Rd, Groton, CT 06340

## Abstract

The lack of a robust gene transformation tool that allows functional testing of the vast number of nuclear genes in dinoflagellates has greatly hampered our understanding of fundamental biology in this ecologically important and evolutionarily unique lineage. Here we report the development of a dinoflagellate expression vector, an electroporation protocol, and successful expression of introduced genes in the dinoflagellate *Oxyrrhis marina*. This protocol, involving the use of Lonza’s Nucleofector and a codon optimized antibiotic resistance gene, has been successfully used to produce consistent results in several independent experiments. It is anticipated that this protocol will be adaptable for other dinoflagellates and will allow characterization of many novel dinoflagellate genes.

## INTRODUCTION

As widely distributed primary producers, essential coral endosymbionts, and the greatest contributors of harmful algal blooms and biotoxins in the ocean, dinoflagellates are a diverse group of unicellular protists with great ecological significance, evolutionary uniqueness, and numerous cytological and genomic peculiarities. Dinoflagellates have immense and permanently condensed genomes with many chromosomes^1–3^; their genomes have a low protein-DNA ratio and histones are functionally replaced with dinoflagellate viral nuclear proteins^4,5^; there are high numbers of repetitive non-coding regions and gene copies, in some species up to ~5,000 copies, organized in tandem arrays^6,7^; only 5-30% of their genes are transcriptionally regulated^7–9^, and microRNAs seem to be the major gene regulating mechanism^10^; and they have undergone extreme plastid evolution, transferring a massive quantity of plastid genes to the nucleus in most of the autotrophic species^11–13^. However, the molecular underpinnings of these unusual features remain elusive. In attempts to address the gap of knowledge, an increasing amount of effort has been made in the last decade to analyze dinoflagellate transcriptomes^14–27^ and genomes^10,28–32^. These experiments have provided not only extensive information on predicted genes and biological pathways, but also an even greater wealth of genes that have weak similarity to characterized proteins or no significant matches in databases. With the increasing volume of dinoflagellate transcriptomic and genomic data, the functional characterization of these novel genes has become a major bottleneck in translating system-level data into a mechanistic understanding of basic dinoflagellate biology, warranting the need for a dinoflagellate genetic transformation system.

Gene transformation attempts have been reported for dinoflagellates by three separate groups. Ten and Miller (1998)^33^ utilized silicon carbide whiskers, polyethylene glycol (PEG), and vigorous shaking to introduce foreign DNA into *Amphidinium* sp. and *Symbiodinium microadriaticum* with a success rate of ~1 ppm. Seventeen years later, Ortiz-Matamoros et al. used PEG, glass beads, shaking and, in some cases, co-incubation with *Agrobacterium tumefaciens* to transform foreign DNA into *Fugacium kawagutii* (formerly *Symbiodinium kawagutii), S. microadriaticum*, and an unclassified *Symbiodiniaceae* species^34,35^. Neither of these reports used codon optimized plasmids for dinoflagellate expression nor did they contain potential dinoflagellate promoters; moreover, both methods remain to be reproduced in other laboratories. In a recent study plasmids containing dinoflagellate minicircle DNA and an antibiotic resist gene were designed and introduced successfully through particle bombardment into the chloroplast genome of the dinoflagellate, *Amphidinium carterae*^36^. Here, we report a successful nuclear gene transformation method for the heterotrophic dinoflagellate, *Oxyrrhis marina*.

*O. marina* is a widespread and ecologically significant heterotrophic dinoflagellate. It is an established model species for both ecological and evolutionary research due its easy cultivable nature, extensive studies related to feeding behavior and nutrition, and its basal position in dinoflagellate phylogeny^37–41^. Although *O. marina* is an early branching dinoflagellate species, it still shares many of the peculiar biological characteristics described above and also retains more typical eukaryotic features that are lacking in later diverging dinoflagellate taxa; thus, it represents a good model for understanding dinoflagellate evolution^41–43^. In addition, *O. marina* has represented planktonic heterotrophs in experiments examining both how they feed and their nutritional value^44–46^. Through various studies as a prey species for copepods and rotifers, *O. marina* has been considered a trophic upgrade as they produce long-chain fatty acids, sterols, and essential amino acids that phytoplankton alone cannot^45,47,48^. Their nutritional value lead to the proposition of using *O. marina* as nutraceuticals for humans and agriculture^45^.

Although *O. marina* lacks a published genome, several transcriptomic studies are available^16,49–52^, and the most exciting finding is *O marina* possess a potential proton pumping rhodopsin with homology to proteorhodopsin^50,51^. Proteorhodopsin is a retinal protein/carotenoid complex that utilizes sunlight to pump protons across a membrane, a non-photosynthetic form of light harvesting^53^. Dinoflagellate species across the phylogenic tree have been found to possess proteorhodopsin homologs, allowing the translational study of this proteins function in *O marina* to the other dinoflagellate species^16,17,50^. Therefore, having a genetic transformation system in place for *O. marina* will greatly excel our understanding of heterotrophic protist ecology, deepen our evolutionary understanding of dinoflagellates within their own branch and relative to other alveolates, allow exploration of the many predicted and novel dinoflagellate genes, and could tap into new industrial applications for *O. marina*, such as a food source or a potential alternative fuel. Additionally, since *O marina* is a heterotrophic species, it is easier to detect the expression of introduced florescent proteins without interference from chlorophyll florescence, as in photoautotrophic species.

In this study, based on genomic and transcriptomic data from several dinoflagellates, we constructed a dinoflagellate expression system (named as DinoIII) that contains potential promoter and termination regions as well as important RNA elements. We incorporated a codon optimized rifampin resistance gene (DinoIII-*arrO*) and green fluorescent protein gene, *gfp*, (DinoIII-*gfp*) into DinoIII, and transformed this DNA as either PCR amplified fragments, excluding the plasmid component for DinoIII-*arrO*, or linear DNA, using a restriction enzyme to digest DinoIII-*gfp*, into *O. marina* using Lonza 4D-Nucleofector^™^ X system (Basel, Switzerland), a gene transformation system enabling transfer of genes directly into the cells’ nucleus^54^. We have been able to repeat transformation for antibiotic resistance several times and verified the presence of both the antibiotic resistance gene and green fluorescent protein several months after transfection.

## RESULTS

### Construction of dinoflagellate backbone expression vector

Initially, the RNA complex sequence from dinoflagellate the *Karenia brevis* (GenBank accession #FJ434727) was inserted into the pMD^™^19-T plasmid vector (Takara, Kusatsu, Shiga Prefecture, Japan) and was used as the vector’s skeleton for a series of modifications. After the addition of more dinoflagellate elements, a functional dinoflagellate backbone vector was achieved, named DinoIII (5137bp; Fig. 1; Supplementary Tables 1-4).

**Fig. 1.**
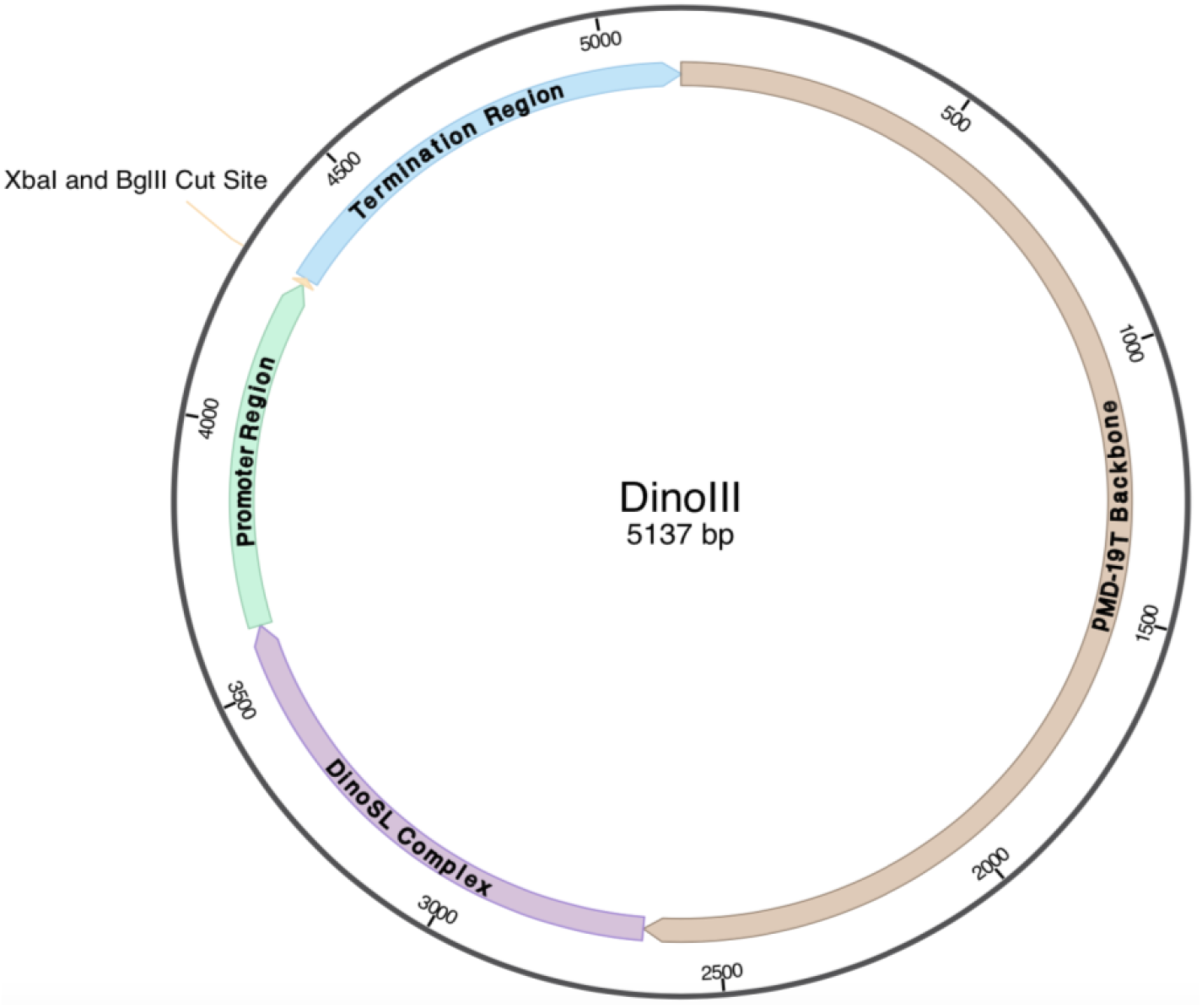
Structure of DinoIII Expression Vector. The bacterial pMD-19 T-Vector portion is in brown, while others depict dinoflagellate elements, including DinoSL Complex region, Promotor Region, and Termination Region. XbaI and BglII Cut site, depicted with a line, that allow for easy gene incorporation in proper orientation.

### Transformation using Lonza 4D-Nucleofector^™^ X Unit system

With Lonza’s 4D-Nucleofector^™^ X Unit, specifically designed for hard-to-transfect cell lines, we went through an extensive cell optimization protocol for *O. marina*, and identified seven adequate pulse code settings. We used these seven pulse codes for follow-up experiments (Table 1). Each pulse code had varying levels of success; for some *gfp* expression was observed, some had a higher percentage of cell survival rate (in which the cells were still growing and dividing after the wildtype NPC cells had died off), whereas others under antibiotic pressure grew to large enough populations to allow for RNA and DNA isolation. Taking all the data into consideration, the pulse codes that showed overall strongest performance were DS-137, DS-134, and DS-120. Nevertheless, we recommend use of all seven settings in the first optimization tests for this species. For other algae to be studied, full optimization tests with the other available solutions should be utilized when using Lonza’s 4D-Nucleofector^™^ X Unit.

**Table 1.**
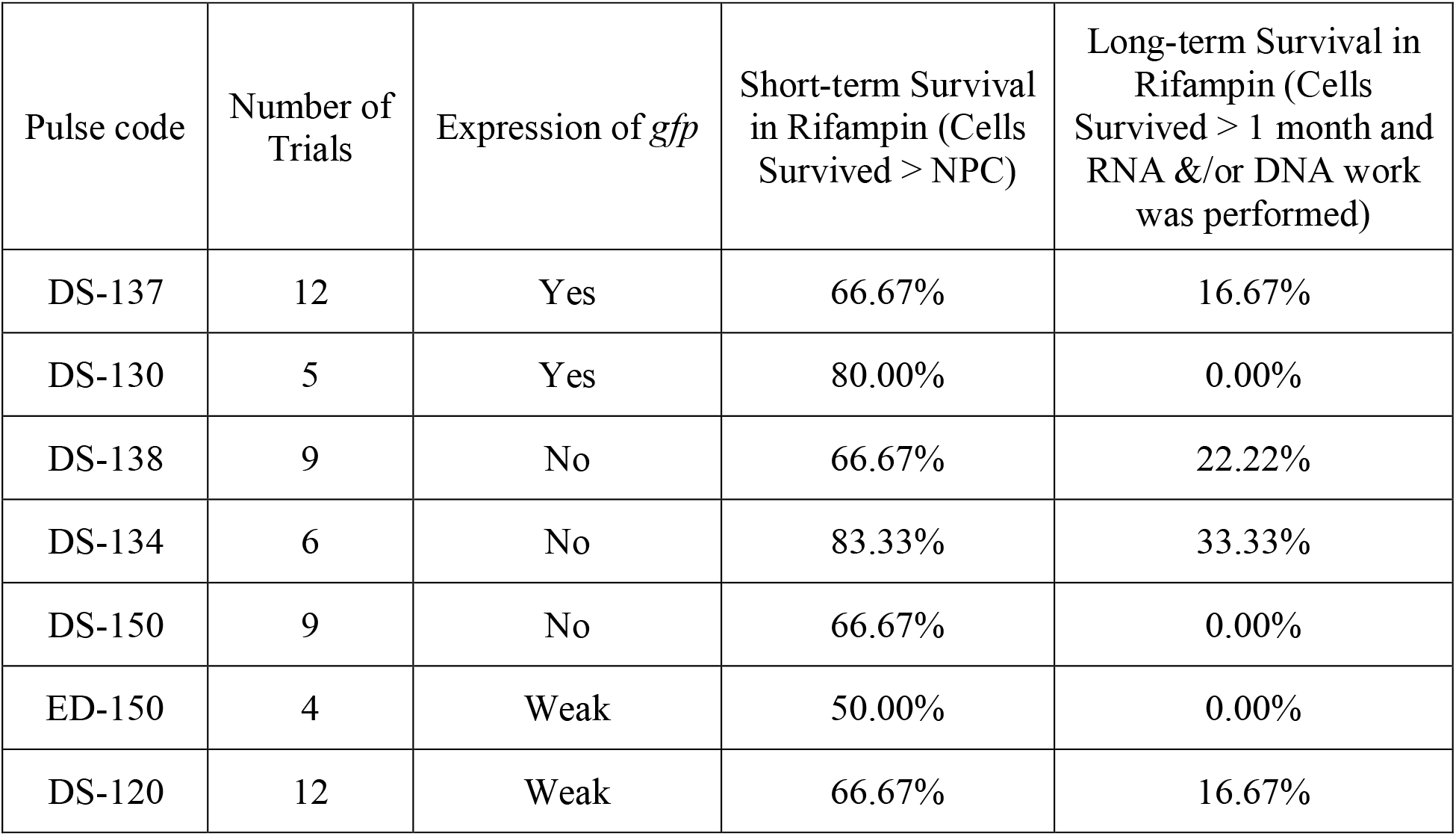
Adequate pulse code settings for transformation of DinoIII-*gfp* and DinoIII-*arrO* into *Oxyrrhis marina*

### GFP Expression

The reporter expression vector, DinoIII-*gfp*, was introduced as linear DNA to *O. marina* cells and the presence of fluorescence was examined microscopically from the third day on. For several weeks we only observed a very dim green signal, but after three months the brightness of the green signal markedly increased for transformed cells using two of the pulse codes, DS-137 and DS-130 (video 1). Perhaps this increase was due to elevated expression after the transformed cells adapted to the new cell environment and/or was the result of accumulating the green fluorescent protein in one single area of the cells (Fig. 2). Less than 1% of the total *O. marina* population in the well was expressing the green fluorescent protein, making the isolation of this cell line challenging without a selection marker.

**Fig. 2.**
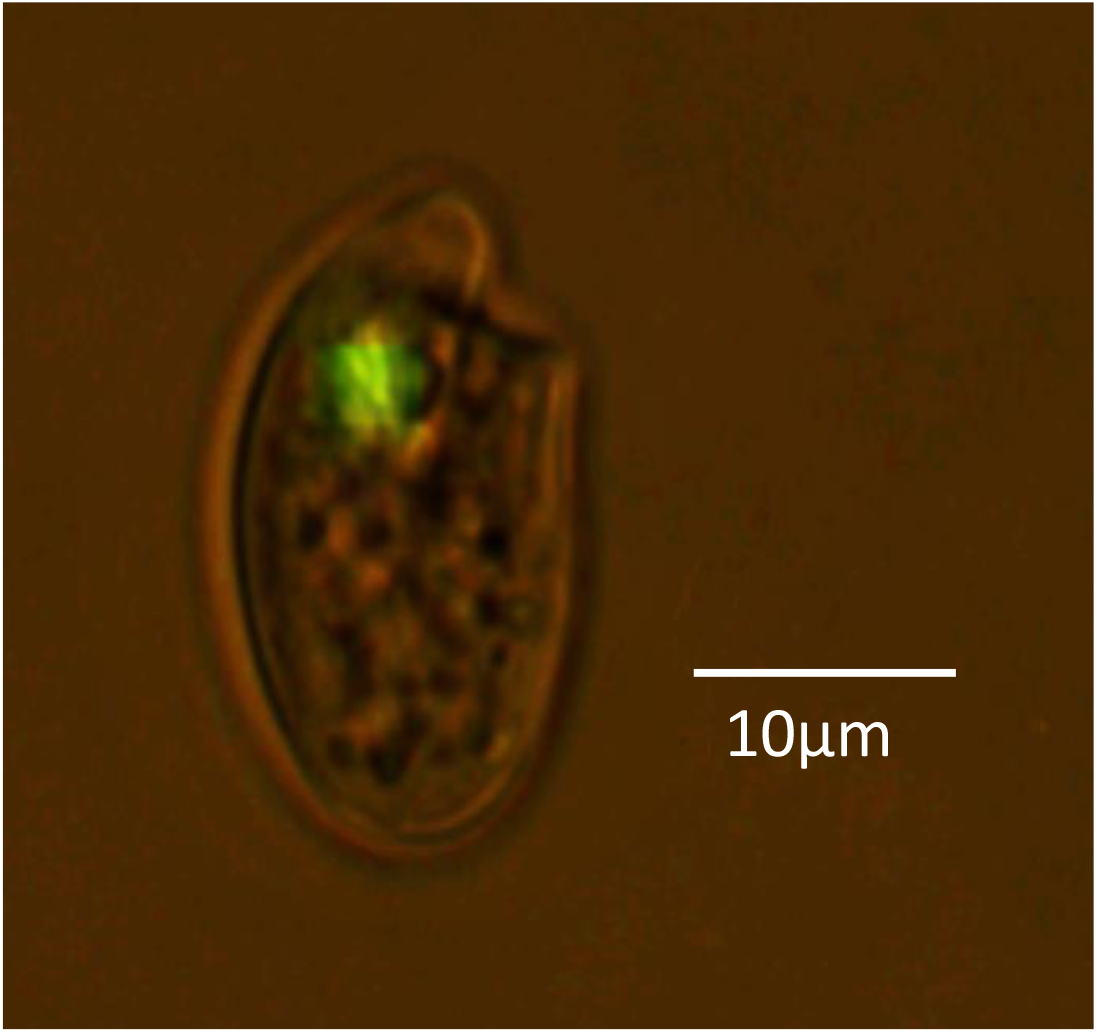
Successful transformation of Oxyrrhis marina with a green florescence protein (GFP) gene (gfp). The green fluorescence indicates expression of the introduced gfp. Photo is a still image of O. marina with a video clip of a *swimming O. marina* cell.

### Rifampin Resistance as a selection marker

To facilitate purification of transformed *O. marina* cells, we screened several selection markers and found rifampin as the most suitable for *O. marina*. Rifampin is an antibiotic used to treat tuberculosis, leprosy, and Legionnaire’s disease and its resistance in bacteria is due to Rifampin ADP-ribosyltransferase activity^55^. We introduced our codon-optimized homolog through our DinoIII vector (DinoIII-*arrO*) and obtained expression of *arrO*, which was verified in several ways. First, the transformed cell culture survived and grew while the wildtype died completely in rifampin-containing medium. Second, after approximately one month we isolated RNA and DNA from both the transformed cells cultured in rifampin-containing medium and a wildtype culture grown in rifampin-free growth medium, and performed reverse-transcription PCR (RT-PCR). We detected the expression of the resistance gene *arrO* only in the experimental treatment and not in the wildtype (Fig. 3A). In addition, we sequenced the PCR product and confirmed that it was *arrO*. Finally, the expression of *arrO* was detected from the cDNA synthesized using Oligo-dT as the primer (Fig. 3B), indicating the transcript of *arrO* was polyadenylated, a phenomenon best known for occurring mostly in eukaryotes mRNA.

Although the cells survived for more than one month, the population increased very slowly and did not seem healthy, probably due to low expression efficiency of the resistance gene. We attempted to increase the expression efficiency of our DinoIII vector by incorporating the intergenic region between *O. marina* rhodopsin tandem repeats, a potential promoter for this highly expressed protein. After introducing the new DinoIII-*arrO*-N PCR fragment into *O. marina* cells we saw an increase in growth rate under antibiotic selection and verified its expression, as reported above, but this time three month after transfection (Fig. 3C). We still, when writing this manuscript, have the cell lines in culture.

**Fig. 3.**
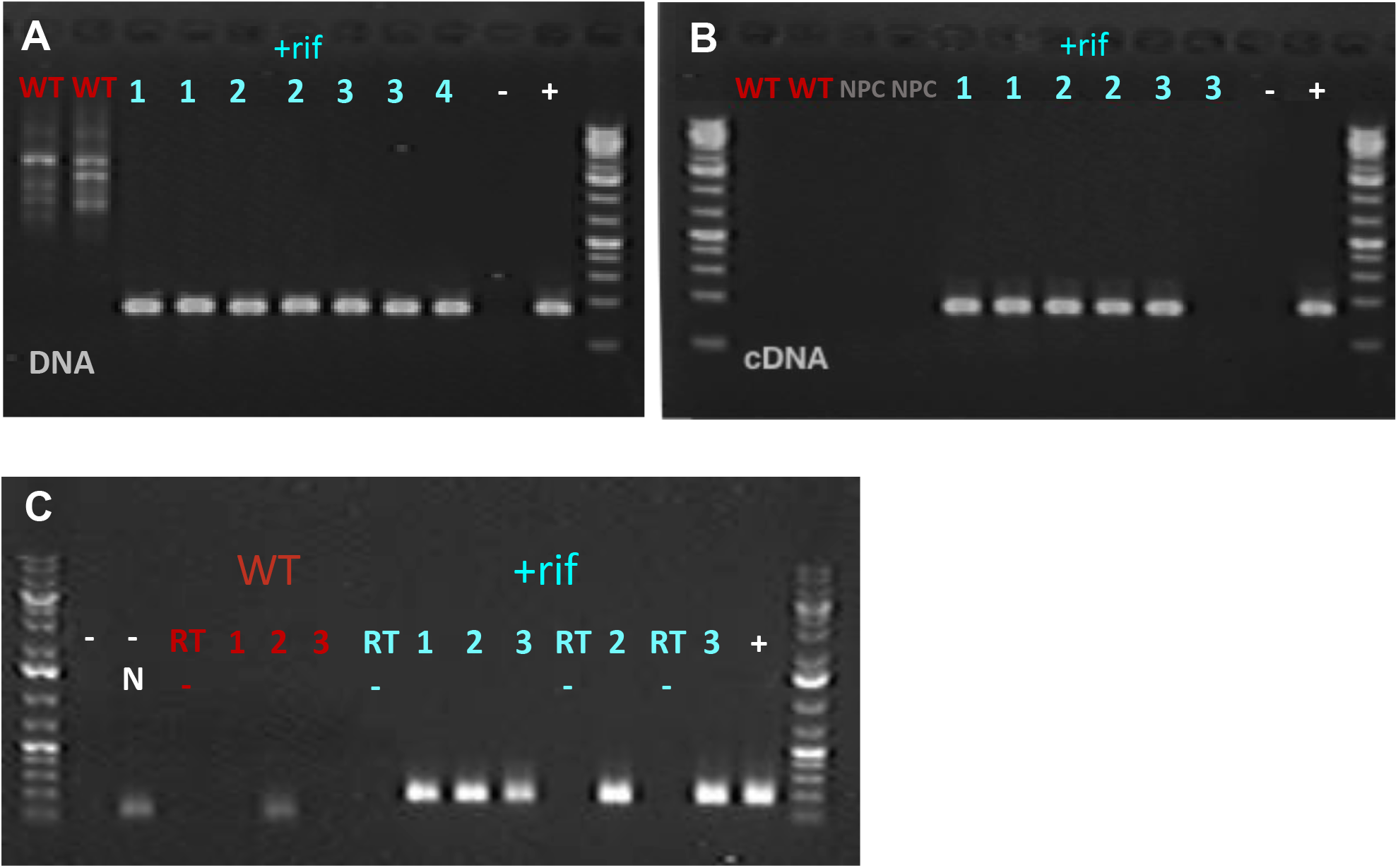
Detection of arrO (rifampin resistance) gene and its expression in the transformed Oxyrrhis marina cells. A) arrO gene detected in genomic DNA of the transformed cells; B) arrO expression detected in the cDNAs of the transformed O. marina. C) arrO-N (arrO plus rhodopsin intergenic region) gene expression detected in the cDNAs of transformed O. marina after 3 months. +rif (in cyano), transformed cells; WT (in red), wild type; “-”, negative control, “-N”, negative nested PCR control; “RT -”, no reverse transcriptase negative control; “1" is N6 library; “2”, OdT Library; “3”, MdT library; “+”, plasmid positive control for arrO-N.

## DISCUSSION

In order to improve understanding of basic dinoflagellate biology, a gene transformation protocol is urgently needed to characterize the function of dinoflagellate genes, particularly the vast number of nuclear genes. A robust and reproducible protocol has been long-awaited. After testing multiple methods (including previously reported ones) and numerous conditions, we have found a passage and herein report a genome-targeted transformation method using a dinoflagellate *gfp* vector (DinoIII-*gfp*) and two dinoflagellate rifampin resistance vectors (DinoIII-*arrO* and DinoIII-*arrO*-N) that were developed based on dinoflagellate genomic and transcriptomic data.

### Effectiveness of the promoter elements

Our efforts began with utilizing expression vectors from the previous reported dinoflagellate transformation^34,35^. This expression system utilized the plant CaMV 35S and *nos* promoters to drive expression of plasmids^34,35^. The *nos* and CaMV 35S promoters have been used extensively for plant transgenic studies, and the CaMV 35S promoter functions in both bacteria and animal systems^56^ as well. These vectors contained the herbicide resistant gene, Basta (glufosinate), as well as several *gfp* fusion genes. Because *O. marina* is not sensitive to Basta, we were looking for green fluorescence as a marker for transformed cells, but it was not observed (results not shown). Therefore, we developed a series of dinoflagellate expression vectors based on existing dinoflagellate transcriptomic and genomic data, mirroring what was previously done for the two model alveolates, *Plasmodium falciparum* and *Tetrahymena thermophile*^57,58^.

In attempts to construct a vector that could drive expression of any gene in any dinoflagellate organism, we included dinoflagellate sequences from two different, phylogenetically separated species, *F. kawagutii* and *K. brevis*. The combination of the included elements (potential promoter, terminator, and RNA elements) were able to drive the expression, albeit at a low level, of the inserted genes in *O. marina*. In order to increase the expression level, we identified a G-rich intergenic region between the highly expressed rhodopsin genes, and incorporated it in our DinoIII*-arrO* vector, yielding visually higher cell survival under antibiotic pressure, indicative of stronger expression of the rifampin resistance gene. When comparing our intergenic region to the luciferase tandem repeats^59^, our sequence is much shorter, only 70 base pairs compared to ~200-2000 and has no real sequence matches in public databanks. No proven promoter exists for dinoflagellates at this time and it is uncertain if the additional sequence contains a promoter. Previous research looking at the binding affinity of *Crypthecodinium cohnii* TATA-binding protein (TBP) homolog and *F. kawagutii* genomic content suggests that dinoflagellates have replaced the typical eukaryotic TATA box with a TTTT motif^10,60^, which is present 65 base pairs upstream from the start codon in our intergenic region and is also present in our “promoter” region with the closest motif 133 base pairs upstream from the start codon. Whether or not these sequences are important can be evaluated in future studies using our method.

### Effectiveness of DNA introduction method

*O. marina* is a naked dinoflagellate that had a difficult time withstanding the physical forces used in previously reported dinoflagellate transformation methods. Electroporation is a gentler method, allowing DNA to pass through temporary pores in an organism’s membrane and has been utilized in many organisms, but requires the removal of seawater and replacement with electroporation buffers, often unable to maintain the osmolality of marine organisms^61^. A new electroporation model, Lonza’s 4D-Nucleofector, provides a user with a score of built-in pulse settings and solutions that remove salts but help maintain dinoflagellates osmolality. The machine has been designed for rapid optimization of both buffer and electric pulse conditions, allowing delivery of nucleic acid substrates into the nucleus^54^. The nucleofector has been widely used on a variety of organisms and cell types and has recently been successfully used on two difficult to transfect marine protists, choanoflagellates and diplonemids^62,63^. For *O. marina*, seven pulse code settings (Table 1) and one solution, SG, allowed the expression of genes in DinoIII vectors. No one pulse code performed the highest across all three of our different criteria. It is interesting to note that DS-138 is considered the weakest pulse setting and DS-120 is the highest of the seven and the remaining five are all in between. Unfortunately, these settings are proprietary and no correlation of the pulse settings can be extracted. Overall, if long-term or short-term expression is the goal, DS-134 is likely the ideal setting.

### Identification of selection marker

Although we were successful in taking microscopic videos of *O. marina* cells expressing the introduced green fluorescent protein, less than 1% of the population actually showed expression. To visualize the green fluorescent protein, cells need to be observed in the dark with blue light, and *O. marina* cells move very fast under the microscope (Video 1), making the isolation of this cell line challenging without a selection marker. The percentage of *O. marina* cells expressing *gfp* decreased with time and the green signal became weaker in four months and eventually was no longer visible. We also discovered that *O. marina* cells would give green-yellowish auto-fluorescence under blue light when fixed with Paraformaldehyde or any other commonly used fixatives, making it very challenging to take a clear image of the *gfp* expressed cell, a necessity to detect the exact location of GFP in the cell (Fig. 2).

Availability of an appropriate selection marker is crucial for yielding a useful transformed cell line. After extensive testing, rifampin was found to be effective for *O. marina*. Rifampin is a very strong pigmented antibiotic that appears to be very light sensitive. Due to this characteristic it is important to keep *O. marina* in lower light settings when under antibiotic selection, keep the cultures well fed, and continue to add new antibiotic medium to the transformed cell lines. Because of the many strains of *O. marina*, it is important to test your species first to determine optimal antibiotic concentrations that can be used to select transformed cell lines, and apply an antibiotic cocktail that will reduce potential microbial communities in the culture.

Ideally a selection marker and a reporter gene can be located on the same plasmid allowing for dual expression. We attempted to put both the *arrO* and *gfp* genes within one single DinoIII vector in multiple arrangements (with or without stop codon in between, fused or not fused) but, unfortunately, the simultaneous expression of both genes was difficult to obtain. Future studies using *arrO* and *gfp* with the rhodopsin intergenic region in between could potentially get over this hurdle.

### Conclusion

Despite the proven challenges, we have developed a dinoflagellate expression system and successfully used it to express foreign genes in the dinoflagellate *Oxyrrhis marina*. Given the extensive studies on *O. marina*, its basal position in the dinoflagellate phylogenetic tree, its easy cultivable nature, and its wide acknowledgement as a model species for heterotrophic protists and dinoflagellates, the gene transformation tool developed here makes the species an even more valuable model. Having a genetic transformation system in place for *O. marina* will allow a deeper understanding of basic dinoflagellate biology. This report is the first stepping block to delve into dinoflagellates molecular biology using *O. marina* as the model and offers a dinoflagellate backbone vector with potential to work across the dinoflagellate phylogenetic tree.

## METHODS

### Culturing *Oxyrrhis marina*

*Oxyrrhis marina* CCMP 1795 was grown at 20°C in autoclaved 0.22mm filtered seawater on a 14:10 hour light:dark cycle at a photon flux of ~100 μ E m^-2^s^-1^ and was fed *Dunaliella tertiolecta* CCMP1320 as prey ^51^. Both species were purchased from the Provasoli-Guillard National Center of Marine Algae and Microbiota in West Boothby Harbor, Maine, USA.

### Constructing Dinoflagellate Expression Vectors

To optimize the utilization of our dinoflagellate expression system several regions were amplified from dinoflagellate genomes and were incorporated to serve as the vector backbone. The first region (974bp) comprises of DNA fragments from the dinoflagellate *Karenia brevis* including SL RNA, SRP RNA, several tRNAs, and U6^64^, which was named as DinoSL complex (Fig. 1; Supplementary Table 1). This region was PCR amplified with DinoSL and KbrSRP-U6R1 primer set (sequences and Tm in Table 2) using the high fidelity PrimeSTAR HS DNA Polymerase (Takara, Kusatsu, Shiga Prefecture, Japan) at 94°C for 1 min, 30 cycles at 95°C for 15s, 58°C for 30s, and 72°C for 1 min, and an additional elongation step at 72°C for 10min. The PCR product was run on 1% agarose to confirm the correct size, purified by passing through a DNA column (Zymo, Irvine, CA, USA), end-fixed, ligated into the pMD^™^19-T plasmid vector (Takara), and transformed chemically into *Escherichia coli* competent cells. Ampicillin was used to select for colonies harboring the region and plasmids were isolated and sequenced to identify the best clone, named as pMD-Dino.

**Table 2.**
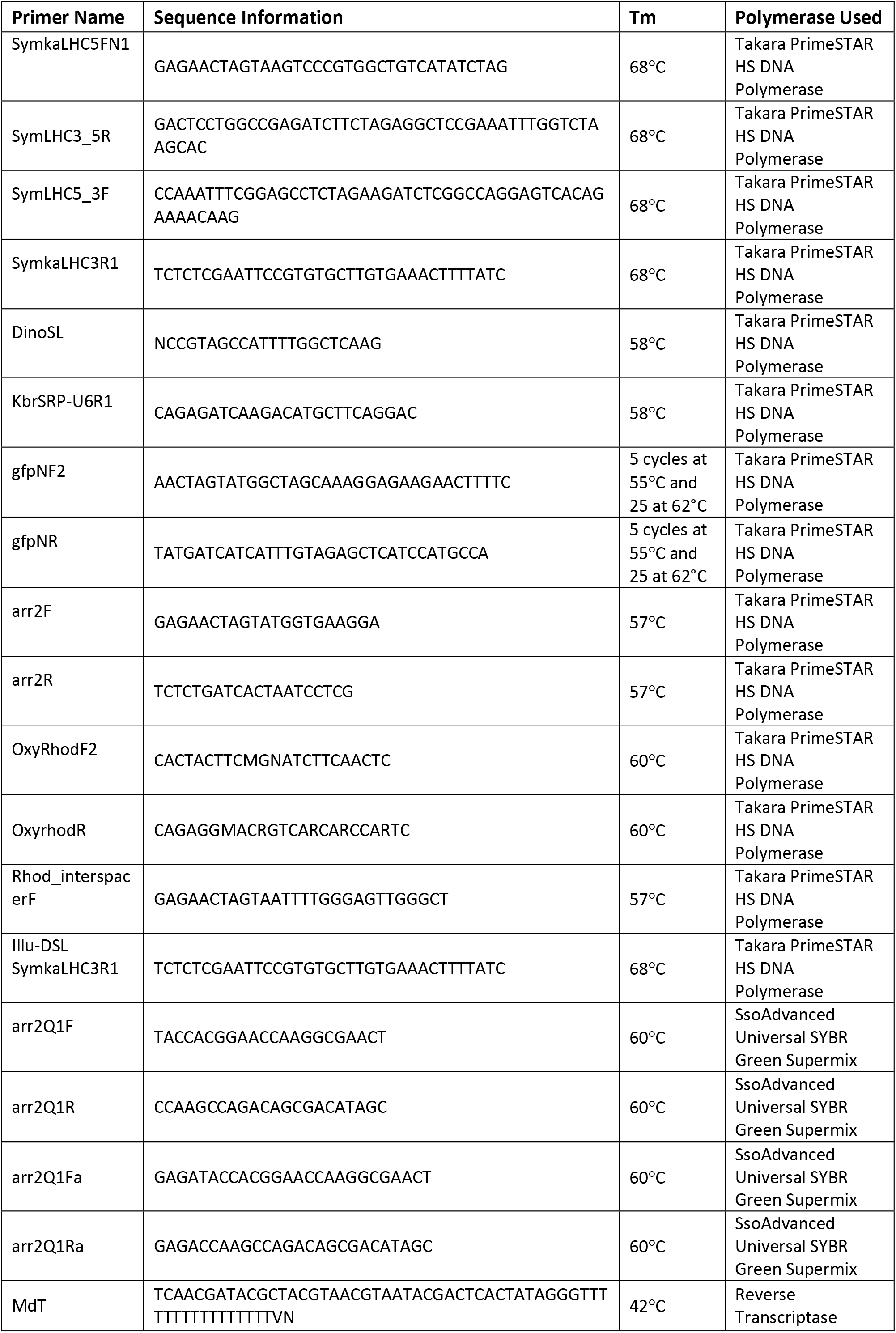
Primers used in the present study

From the *F. kawagutii* genome sequence data^10^, we located the highly expressed light harvesting complex (LHC) gene. Its upstream “promoter” region (672bp; Supplementary Table 2) and downstream “termination” region (812bp; Supplementary Table 3) were PCR-amplified using the following primer sets: SymkaLHC5FN1 and SymLHC3_5R for the “promoter” and SymLHC5_3F and SymkaLHC3R1 for the “termination” region. All PCRs were performed at 94°C for 1 min, 25 cycles at 95°C for 15s, 68°C for 30s, and 72°C for 1 min, and 1 cycle of 72°C for 10 min. The sizes of the amplicons were checked by electrophoresis and DNA was purified by passing through a DNA column (Zymo).

SymLHC3_5R and SymLHC5_3F were designed to contain an overhang of either a portion of the “termination” region or the “promoter” region, respectfully, in order to link the two PCR products; thus, the two products were used in an equal molar ratio as template for the second PCR at 94°C for 1 min, 5 cycles without primers at 95°C for 15s, 68°C for 90s, 20 cycles with SymkaLHC5FN1 and SymkaLHC3R1 at 95°C for 15s, 68°C for 90s, and an extra elongation step of 72°C for 10 mins. The single product was checked by electrophoresis to verify the amplicon size, gel isolated, and digested with SpeI and EcoRI as SymkaLHC5FN1 had a SpeI site and SymkaLHC3R1 had an EcoRI site added to their 5’ ends for easy incorporation into pMD-Dino harboring the SL RNA, SRP RNA, several tRNAs, and U6 region. The pMD-Dino vector was digested with XbaI and EcoRI and treated with alkaline phosphatase to avoid self-ligation. After 3.5 hours of digestion both products were purified by ethanol precipitation and ligated overnight in a 2:1 molar ratio (LHC product:vector) and transformed into competent *E. coli* cells. The colonies obtained were picked randomly, and plasmids were isolated and sequenced to identify the best clone harboring the correct DinoSL Complex-LHC region sequence, giving rise to the dinoflagellate expression vector backbone, DinoIII (5137bp; Fig. 1; Supplementary Table 4).

SymLHC3_5R and SymLHC5_3F primers were designed to also have an XbaI and BglII site in between the “Promoter” and “termination” regions so that a gene, either a reporter or an antibiotic resistant gene, could be inserted in the correct orientation. Accordingly, both a *gfp* gene and a rifampin resistance gene were incorporated into the expression vector, to yield DinoIII*-gfp* and DinoIII-*arrO*, respectively (Supplementary Table 5,6). For DinoIII-*gfp*, the crystal jelly *Aequorea victoria gfp* was amplified from the pGlo^™^ Plasmid (Bio-Rad, Hercules, CA, USA) using gfpNF2 and gfpNR at 94°C for 1 min, 5 cycles at 95°C for 15s, 55°C for 30s, and 72°C for 30s, 25 cycles at 95°C for 15s, 62°C for 30s, and 72°C for 30s, and an extra elongation step of 72°C for 10 mins. For *arrO*, a homolog to Rifampin ADP-ribosylating transferase from bacterium *Citrobacter freundii* was found on GenBank (accession # NC_019991) and was codon-optimized for *O. marina* based on codon usage data from reported *O. marina* genes, and was synthesized through GeneArt (ThermoFisher Scientific, Waltham, MA, USA). Upon arrival the synthesized *arrO* was PCR amplified at 94°C for 1 min, 30 cycles at 95°C for 15s, 57°C for 30s, and 72°C for 30s, and an extra elongation step of 72°C for 10 mins with arr2F and arr2R primers. Both the *gfp* and *arrO* genes had a SpeI site at 5’-end and a BclI site at 3’-end; thus, after their PCR amplification, the single products were checked on a gel, passed through DNA columns to purify, and digested with SpeI and BclI for 3.5 hours. At the same time DinoIII was digested with XbaI and BglII and treated with alkaline phosphates. After digestion, the *gfp* and *arrO* genes were ligated into DinoIII overnight in a 2:1 molar ratio, and were transformed into competent *E. coli* cells. Plasmids were isolated and sequenced to identify the clones containing correct sequences of DinoIII*-gfp* and DinoIII-*arrO*, respectively.

### Optimizing promoter region

To optimize the expression of transformed genes for *O. marina*, we set out to find a promoter region for their highly expressed proteorhodopsin genes (2-4 X10^6^ copies/ng total RNA, twice that of mitochondrial *cox1*)^51^. To do this we used the OxyRhodF2 and OxyrhodR primer set^51^, under thermal cycle conditions with an extended extension time to favor long amplicons that cover two or more tandem repeats of the gene: 94°C for 1 min, 25 cycles at 95 °C for 15 s, 60°C for 30s, and 72°C for 90s, and an extra elongation step of 72°C for 10 min. Bands of ~800bp and ~1400bp were gel purified, cloned, and sequenced to identify a potential promoter. This yielded an intergenic region between rhodopsin tandem repeats, with the following sequence: aattttgggagttgggctggaagatggggttggtggggatcgggggagaggtgactggtgtgtggtcgag. We added this sequence to the 5’-end of the *arrO* gene through GeneArt (ThermoFisher Scientific) and incorporated this *arrO*-N sequence into the DinoIII vector (DinoIII-*arrO*-N; Supplementary Table 7) as described above.

### Introducing DNA into *O. marina* using Lonza’s Nucleofector

*O. marina* cultures were fed with *D. tertiolecta* three days before transformation in order to reach high cell densities. Taking advantage of the species photo-tactic behavior, *O. marina* cells were concentrated using a flashlight, allowing cells to swim toward the light, consequently gathering only healthy cells from the culture. The cultures cell numbers were counted microscopically using a Sedgwick-Rafter counting chamber.

Electroporation was carried out using Lonza 4D-Nucleofector^™^ X Unit system in 16-well Nucleocuvette^™^ Strips using the manufacturer’s SG and Supplemental 1 solutions. DinoIII-*gfp* was digested with EcoRI and introduced as a liner plasmid and DinoIII-*arrO* plasmid was PCR amplified, to produce linear fragments that only contained the dinoflagellate DNA portion. PCR was carried out using Illu-DSL and SymkaLHC3R1 primers at 94°C for 1 min, 25 cycles at 95°C for 15s, 68°C for 90s, and an extra elongation step of 72 °C for 10 mins. The linear plasmid and PCR product were checked through electrophoresis, retrieved, and concentrated to 1 μg/μl using a Millipore Microcon DNA Fast Flow Column (Burlington, MA, USA). For each transformation well, 16.4 μl of solution SG, 3.6 μl of Supplemental 1 solution, and 2 μl of PCR product were used as transformation solution.

For every well, ca. 2.5 × 10^5^ cells were added and all the cells for the experiment (including the controls) were collected in 50mL tubes. The cells were centrifuged at 2500g for 3 mins, enough to form a pellet at the bottom of the tubes, and all but ~2-3mL of medium was removed. The cells were transferred into 1.5mL tubes and centrifuged at 900g for 2 mins and all remaining liquid was removed. The cells were re-suspended in the transformation solution and 22 μl were added to each well. After an initial optimization test the following electroporation settings were used for further experiments: DS-137, DS-130, DS-138, DS-134, DS-150, ED-150, DS-120, and no pulse controls (NPC).

Immediately after electroporation 80 μl of the same seawater medium (SW) that *O. marina* was cultured on but with an antibiotic cocktail, AKS (100 μg/μl ampicillin, 50 μg/μl kanamycin, and 50 μg/μl streptomycin), was added to each well. All of the volume was gently transferred into 24-well plates where each well already contained 1.4mL of the same SW+AKS medium. The transformed cells were allowed to recover for three days. For the DinoIII-*gfp* transformations, cells were examined microscopically under blue light for *gfp* expression. For DinoIII-*arrO* cells, 750 μl were transferred to new 24-well plates and 750 μl SW+AKS containing 450 μg/μl of rifampin was added to both plates, so that the final concentration of rifampin was 225 μg/μl. On the third day under antibiotic selection, *D. tertiolecta* in 225 μg/μL rifampin medium was added to each well. New antibiotic solution was added every 3 weeks but the concentration was dropped down to 200 μg/μL to allow for greater cell growth and *D. tertiolecta* in 225 μg/μL rifampin medium was supplied whenever they were no longer detected in the medium.

### Detecting the transformed gene and its expression

Total DNA was isolated from both wildtype (WT) and +*arrO O. marina* cultures using our CTAB method^65^ and total RNA was isolated using the Trizol-Chloroform method in combination with Zymo Quick RNA Miniprep Kit (Irvine, CA, USA)^66^. As the transformed cultures grew slowly under antibiotic pressure, only 200-400 cells were available. These cells were divided into two for DNA and collected on TSTP Isopore 3μm membrane filters (MilliporeSigma, Burlington, MA, USA). DNA and RNA were isolated as reported^16,66^.

Several first-strand cDNA preparations were made with RNA isolated using ImProm-II^™^ Reverse Transcriptase (Promega, Madison, WI, USA) following the manufacture’s protocol with random hexamer (N6), oligo(dT)18 (OdT), and modified OdT (MdT; Table 2) as the primers, respectively. If cell numbers were very low only OdT was used for cDNA synthesis to maximize cDNA production. Negative controls were included where no reverse transcriptase was added and was instead replaced with DEPC water.

PCR was performed using both the DNA and cDNA as templates with primer set arr2Q1Fa-arr2Q1Ra. Due to the low number of transformed cells used in DNA and RNA isolation, this PCR did not yield detectable amounts of products. The PCR products were then diluted 1000- and 10000-fold and used as template for a nested PCR with arr2Q1F-arr2Q1R as the primer (Table 2) for quantitative PCR. The products were run on a gel and sequenced directly to ensure the *arrO* gene was correctly amplified.

## Acknowledgements

We thank Dr. Lu Wang for her help in constructing DinoIII and Sarah Baseler for her assistance in screening selection markers. The work was supported by the Gordon and Betty Moore Foundation through Grant GBMF4980 (to SL and HZ) and the National Science Foundation Graduate Research Fellowship under Grant No. 1247393 (to BNS). Any opinions, findings, and conclusions or recommendations expressed in this material are those of the authors and do not necessarily reflect the views of the National Science Foundation.

---

Video 1. The *Oxyrrhis marina* cell transformed with *gfp* showing green florescence under blue light. Video was taking using Olympus BX51 microscopic system with a DP74 Olympus camera under 100X magnification.

**Video in Supplementary Material**

