## Supplemental Table for "Nuclear gene transformation in a dinoflagellate"

Supplementary Table. 1 The RNA complex sequence from dinoflagellate *Karenia brevis* (GenBank accession # FJ434727) containing SL RNA, SRP RNA, several tRNAs, and U6. (Sequence yellow highlighted).

| Name | Sequence |
|---|---|
| RNA Elements | CCCGTAGCCATTTTGGCTCAAGGTACAAGTCGGGCTGATGCGGTCACGCAGG<br>CCTCTTTTGTATCAGATAAGACAGCAGACGTACATGATATAAATATTTATT<br>ATCGCGATGCTCCATCTGCAAAATTCCACTTGGCAGAGAAGATTTTCAATG<br>CATGAAAACCTCGCAGAACCAAGCACCAATGTCAGATCATGTATGAGTCCT<br>AGGCGCCAGTAAGTCTTTCAATTGGGTGATGGCCACATTACCTGTTCTCTTC<br>CTGAGGGATGCAGAATGTGGTGTGAGTGGTTGCCGAATGCTGTGCTGAACA<br>TGTAGATGGGCTGCCCCAGCCAAGTGGATAACCTACAAGGACTGCAGGATG<br>TAGATCCAGCAGTCCTGGTAGATAATGATTCAGACACCGGCTCAGGCTGGC<br>AACAGAGCAGGCAAACCTGAACACTATCGCGCAGTGGTGGGAATACAGCCC<br>AGAATGGGAAGCAACCTGGCCTCTTCAGCCAGCCATGACCACAAGCAACTCT<br>GCAGTTTTGCAACACGAAATCTTTCAGGGTCTGCTCCCTGTGCTTACACATA<br>TTCAAAGATATTCTTCTGCGCATTTCAATGCAAATTCTTGGCGATGATTCG<br>CTTCCGCAGGTATAGTGTTACTAGTATATAAGTCTAATCATATACTTCAAT<br>TTATTAGTAATATTGTGAATTTCTAGGCAAGAAGGATGCCTGGACTCTGAA<br>TTATTATGGCATTGAGTAGAATCTGGATCTTGATGATTATGCATTAATATC<br>TTGAAATGCATTTGGATTCCCTTCGGGGATCATCCGTTAAAATTGGAACGA<br>TACAGAGAAGATTAGCATGGCCCCTGCGCAAGGATGACACGCACAAATCGA<br>GAAGTGTAAACAATTTTTTTGAAATTAATTGCCACTTTATTTTGAATACCT<br>GAATATGCAGGTGAAGTAGTATAAGGTATTCATTATCATTGCGATTTTCTA<br>G |

Supplementary Table 2. The "promoter" region sequence containing the upstream region of the highly expressed light harvesting complex (LHC) gene of dinoflagellate *Fugacium kawagutii* (formerly *Symbiodinium kawagutii*). (Sequence blue highlighted).

| Name | Sequence |
|---|---|
| Promoter Region | TCCCGTGGCTGTCATATCTAGTAACCCTCACCTGGCAGGTGGGGAAAAGGC<br>GAAGACAAATAGAATCAAATAGAATGTATGTGCTGACGTAGGGCTACATTA<br>CCTGAGGCTTGAGGAACCTGTGGGTATGAATGCTACTGTTGGCAGCATCCA<br>GTAGGTTCAGAAACACACGTCCCCACGTAATTTTTTGCATGTCTAAATGCA<br>CTGTGATGATTGATGATGATTGATTCCTTGAAGGCCATGGTGGGTTACTCT<br>GTGGCCTCTTGCCACTGACTTCCAGGAAAACGCGGATTTTCTGACCATCACA<br>AGGGACCACTACGAGAAGTATGCAGCGATCTCAGCAGAGTTCATTTACTGA<br>CCATTCTCAGGAAGATCGCTCGGGATCACTTGGGTAATATTGCTTTCTTCGC<br>AGATGTTTTTTCCCTATTAAGCTTTTTGAGTCCCCTCATCGAGTGCCGCATA<br>AGTTTTTGCGTTGACATTGGCGGCAAAAAGTAAGACTAAACGATAGTTGCT<br>TCAAGCACGCTTCTCAATCAACATTTTTTCAAATCAAATGGGCACAGGCAG<br>GTACCATCCAGTCCCGCAGAAGTGCTTATATATATATATATTATACACTCA<br>CGTGAGTGTGTTCAATGCAGGTCTGTTCCTCAAGTGCTTAGACCAAATTTC<br>GGAGCC |

Supplementary Table 3. The “termination” region containing the downstream sequence of the highly expressed light harvesting complex (LHC) gene of dinoflagellate *Fugacium kawagutii* (formerly *Symbiodinium kawagutii*). (Sequence pink highlighted).

| Name | Sequence |
|---|---|
| Termination Region | CGGCCAGGAGTCACAGAAAACAAGATCACTTGGAGATGTTTCAATCCCGAC<br>TTGTGTCGTGCCAGAGTGCTACTTGAAAACTTGAAAATTGCGGACTGTCAT<br>GGATTCGCCCTTGTCTTGTGATCCTTTTTTTGGGGGAGCCAGGTGAGAACA<br>ATGTTGTCGATGTGCTTATTTGGCTTCGCAGTCAAAACATGGGATACTTGA<br>GACATGAAAGAAAAATGCCGCAACGATAGCTCCATCCAATTCCATTCAGCT<br>CCGACTACAGATGATAGCGCTTGACACCAATGACATGCTTGTACAGCTGCC<br>ATTTGGAAGGCAGGGAAGCTCCATAAGCTCGGGTCCCCAGGACTTTGGTCG<br>GTCTCACATCAGATTCGGCTAGCCAGCCCATAGCAGCCGCGGGAGATTTCGG<br>TTGTTTGCTACAATGATTGGGGCGCCTTTCTGCGAACTTTGTGACATGTTTC<br>CTCAAAATGTCAAGCAATTTTGATCTTAAAAGTTTTGATAATGCTTGCTTC<br>CACAAGCGACCTACAGTAGGAAATGTCTCCACAATCTCCACAGATTCAGGA<br>CTCATCACTATGTGTGCCGTGCAGGGGTAGGGCGCAGACATGACAACATAC<br>AACACACATGAACTAAAGAATCCAAGTCGCGGACAAAAAAATCTGATCTTA<br>CACTTACACAGAATGCAGGTTATTAGCGACGCTTCCATTGCCACCGGAGTG<br>GCAATCGTTGAGGCGCTTCATCGAACAGAGGGTGAACTTCTTGAGGCTGGG<br>AGGACCGCGCAGATGCGGCTGATAAAAGTTTCACAAGCACACGGA |

Supplementary Table 4. Backbone of Dinoflagellate expression vector, containing pMD-19 T-Vector (shown in lowercase) and the RNA Elements (yellow highlighted), Promoter Region (blue highlighted), and Termination Region (pink highlighted). The restriction enzyme sites for XbaI (**tctaga**) and BglII (**agatct**) shown in bold are induced for cloning the gene of interest in the correct orientation.

| Name | Sequence |
|---|---|
| DinoIII | gacgaaagggcctcgtgatacgcctatttttataggttaatgtcatgataataatggtttcttagacgtca<br>ggtggcacttttcggggaaatgtgcgcggaacccctatttgtttatttttctaaatacattcaaatatgtat<br>ccgctcatgagacaataaccctgataaatgcttcaataatattgaaaaaggaagagtatgagtattcaa<br>catttccgtgtcgcccttattcccttttttgcggcattttgccttcctgtttttgctcacccagaaacgctggt<br>gaaagtaaaagatgctgaagatcagttgggtgcacgagtgggttacatcgaactggatctcaacagcg<br>gtaagatccttgagagttttcgccccgaagaacgttttccaatgatgagcacttttaaagttctgctatgtg<br>gcgcggtattatcccgtattgacgccgggcaagagcaactcggtcgccgcatacactattctcagaatg<br>acttggttgagtactcaccagtcacagaaaagcatcttacggatggcatgacagtaagagaattatgca<br>gtgctgccataaccatgagtgataacactgcggccaacttacttctgacaacgatcggaggaccgaagg<br>agctaaccgcttttttgcacaacatgggggatcatgtaactcgccttgatcgttgggaaccggagctgaa<br>tgaagccataccaaacgacgagcgtgacaccacgatgcctgtagcaatggcaacaacgttgcgcaaac<br>tattaactggcgaactacttactctagcttcccggcaacaattaatagactggatggaggcggataaagt<br>tgcaggaccacttctgcgctcggcccttccggctggctggtttattgctgataaatctggagccggtgagc<br>gtgggtctcgcggtatcattgcagcactggggccagatggtaagccctcccgtatcgtagttatctacac<br>gacggggagtcaggcaactatggatgaacgaaatagacagatcgctgagataggtgcctcactgatta<br>agcattggtaactgtcagaccaagtttactcatatatactttagattgatttaaaacttcatttttaattta

| | |
|---|---|
| | aaggatctaggtgaagatcctttttgataatctcatgaccaaaatcccttaacgtgagttttcgttccactg<br>agcgtcagaccccgtagaaaagatcaaaggatcttcttgagatcctttttttctgcgcgtaatctgctgctt<br>gcaaacaaaaaaaccaccgctaccagcggtggtttgtttgccggatcaagagctaccaactctttttccg<br>aaggtaactggcttcagcagagcgcagataccaaatactgttcttctagtgtagccgtagttaggccacc<br>acttcaagaactctgtagcaccgcctacatacctcgctctgctaatcctgttaccagtggctgctgccagt<br>ggcgataagtcgtgtcttaccgggttggactcaagacgatagttaccggataaggcgcagcggtcggg<br>ctgaacggggggttcgtgcacacagcccagcttggagcgaacgacctacaccgaactgagatacctac<br>agcgtgagctatgagaaagcgccacgcttcccgaagggagaaaggcggacaggtatccggtaagcgg<br>cagggtcggaacaggagagcgcacgagggagcttccagggggaaacgcctggtatctttatagtcctg<br>tcgggtttcgccacctctgacttgagcgtcgatttttgtgatgctcgtcaggggggcggagcctatggaa<br>aaacgccagcaacgcggcctttttacggttcctggccttttgctggccttttgctcacatgttctttcctgcg<br>ttatcccctgattctgtggataaccgtattaccgcctttgagtgagctgataccgctcgccgcagccgaac<br>gaccgagcgcagcgagtcagtgagcgaggaagcggaagagcgcccaatacgcaaaccgcctctcccc<br>gcgcgttggccgattcattaatgcagctggcacgacaggtttcccgactggaaagcgggcagtgagcg<br>caacgcaattaatgtgagttagctcactcattaggcaccccaggctttacactttatgcttccggctcgtat<br>gttgtgtggaattgtgagcggataacaatttcacacaggaaacagctatgaccatgattacgccaagctt<br>gcatgcctgcaggtcgacgattCCCGTAGCCATTTTGGCTCAAGGTACAAGTCGGGC<br>TGATGCGGTCACGCAGGCCTCTTTTGTATCAGATAAGACAGCAGACGTACA<br>TGATATAAATATTTATTATCGCGATGCTCCATCTGCAAAATTCCACTTGGC<br>AGAGAAGATTTTCAATGCATGAAAACCTCGCAGAACCAAGCACCAATGTCA<br>GATCATGTATGAGTCCTAGGCGCCAGTAAGTCTTTCAATTGGGTGATGGCC<br>ACATTACCTGTTCTCTTCCTGAGGGATGCAGAATGTGGTGTGAGTGGTTGC<br>CGAATGCTGTGCTGAACATGTAGATGGGCTGCCCCAGCCAAGTGGATAACC<br>TACAAGGACTGCAGGATGTAGATCCAGCAGTCCTGGTAGATAATGATTCAG<br>ACACCGGCTCAGGCTGGCAACAGAGCAGGCAAACCTGAACACTATCGCGCA<br>GTGGTGGGAATACAGCCCAGAATGGGAAGCAACCTGGCCTCTTCAGCCAGC<br>CATGACCACAAGCAACTCTGCAGTTTTGCAACACGAAATCTTTCAGGGTCT<br>GCTCCCTGTGCTTACACATATTCAAAGATATTCTTCTGCGCATTTCAATGCA<br>AATTCTTGGCGATGATTCGCTTCCGCAGGTATAGTGTTACTAGTATATAAG<br>TCTAATCATATACTTCAATTTATTAGTAATATTGTGAATTTCTAGGCAAGA<br>AGGATGCCTGGACTCTGAATTATTATGGCATTGAGTAGAATCTGGATCTTG<br>ATGATTATGCATTAATATCTTGAAATGCATTTGGATTCCCTTCGGGGATCA<br>TCCGTTAAAATTGGAACGATACAGAGAAGATTAGCATGGCCCCTGCGCAAG<br>GATGACACGCACAAATCGAGAAGTGTAAACAATTTTTTTGAAATTAATTGC<br>CACTTTATTTTGAATACCTGAATATGCAGGTGAAGTAGTATAAGGTATTCA<br>TTATCATTGCGATTTTCTAGTAAGTCCCGTGGCTGTCATATCTAGTAACCCT<br>CACCTGGCAGGTGGGGAAAAGGCGAAGACAAATAGAATCAAATAGAATGTA<br>TGTGCTGACGTAGGGCTACATTACCTGAGGCTTGAGGAACCTGTGGGTATG<br>AATGCTACTGTTGGCAGCATCCAGTAGGTTCAGAAACACACGTCCCCACGT<br>AATTTTTGCATGTCTAAATGCACTGTGATGATTGATGATGATTGATTCCT<br>TGAAGGCCATGGTGGGTTACTCTGTGGCCTCTTGCCACTGACTTCCAGGAAA<br>ACGCGGATTTTCTGACCATCACAAGGGACCACTACGAGAAGTATGCAGCGA<br>TCTCAGCAGAGTTCATTTACTGACCATTCTCAGGAAGATCGCTCGGGATCAC<br>TTGGGTAATATTGCTTTCTTCGCAGATGTTTTTTCCCTATTAAGCTTTTTG<br>AGTCCCCTCATCGAGTGCCGCATAAGTTTTTGCGTTGACATTGGCGGCAAA |

| | |
|---|---|
| | AAGTAAGACTAAACGATAGTTGCTTCAAGCACGCTTCTCAATCAACATTTT<br>TTCAAATCAAATGGGCACAGGCAGGTACCATCCAGTCCCGCAGAAGTGCTT<br>ATATATATATATATTATACACTCACGTGAGTGTGTTCAATGCAGGTCTGTT<br>CCTCAAGTGCTTAGACCAAATTTCGGAGCC**tctagaagatct**CGGCCAGGA<br>GTCACAGAAAACAAGATCACTTGGAGATGTTTCAATCCCGACTTGTGTCGT<br>GCCAGAGTGCTACTTGAAAACTTGAAAATTGCGGACTGTCATGGATTCGCC<br>CTTGTCTTGTGATCCTTTTTTTGGGGGAGCCAGGTGAGAACAATGTTGTCG<br>ATGTGCTTATTTGGCTTCGCAGTCAAAACATGGGATACTTGAGACATGAAA<br>GAAAAATGCCGCAACGATAGCTCCATCCAATTCCATTCAGCTCCGACTACAG<br>ATGATAGCGCTTGACACCAATGACATGCTTGTACAGCTGCCATTTGGAAGG<br>CAGGGAAGCTCCATAAGCTCGGGTCCCCAGGACTTTGGTCGGTCTCACATCA<br>GATTCGGCTAGCCAGCCCATAGCAGCCGCGGGAGATTTCGGTTGTTTGCTAC<br>AATGATTGGGGCGCCTTTCTGCGAACTTTGTGACATGTTTCCTCAAAATGT<br>CAAGCAATTTTGATCTTAAAAGTTTTGATAATGCTTGCTTCCACAAGCGAC<br>CTACAGTAGGAAATGTCTCCACAATCTCCACAGATTCAGGACTCATCACTAT<br>GTGTGCCGTGCAGGGGTAGGGCGCAGACATGACAACATACAACACACATGA<br>ACTAAAGAATCCAAGTCGCGGACAAAAAAATCTGATCTTACACTTACACAG<br>AATGCAGGTTATTAGCGACGCTTCCATTGCCACCGGAGTGGCAATCGTTGA<br>GGCGCTTCATCGAACAGAGGGTGAACTTCTTGAGGCTGGGAGGACCGCGCA<br>GATGCGGCTGATAAAAGTTTCACAAGCACACGGAgaattcactggccgtcgttttaca<br>acgtcgtgactgggaaaaccctggcgttacccaacttaatcgccttgcagcacatccccctttcgccagc<br>tggcgtaatagcgaagaggcccgcaccgatcgcccttcccaacagttgcgcagcctgaatggcgaatg<br>gcgcctgatgcggtattttctccttacgcatctgtgcggtatttcacaccgcatatggtgcactctcagtac<br>aatctgctctgatgccgcatagttaagccagccccgacacccgccaacacccgctgacgcgccctgacg<br>ggcttgtctgctcccggcatccgcttacagacaagctgtgaccgtctccgggagctgcatgtgtcagagg<br>ttttcaccgtcatcaccgaaacgcgcga |

Supplementary Table 5: Sequence for a homolog to crystal jelly *Aequorea victoria gfp* obtained from the pGlo™ Plasmid.

| Name | Sequence |
|---|---|
| *gfp* | ATGGCTAGCAAAGGAGAAGAACTTTTCACTGGAGTTGTCCCAATTCTTGTT<br>GAATTAGATGGTGATGTTAATGGGCACAAATTTTCTGTCAGTGGAGAGGGT<br>GAAGGTGATGCTACATACGGAAAGCTTACCCTTAAATTTATTTGCACTACT<br>GGAAAACTACCTGTTCCATGGCCAACACTTGTCACTACTTTCTCTTATGGTG<br>TTCAATGCTTTTCCCGTTATCCGGATCATATGAAACGGCATGACTTTTTCAA<br>GAGTGCCATGCCCGAAGGTTATGTACAGGAACGCACTATATCTTTCAAAGA<br>TGACGGGAACTACAAGACGCGTGCTGAAGTCAAGTTTGAAGGTGATACCCT<br>TGTTAATCGTATCGAGTTAAAAGGTATTGATTTTAAAGAAGATGGAAACAT<br>TCTCGGACACAAACTCGAGTACAACTATAACTCACACAATGTATACATCAC<br>GGCAGACAAACAAAAGAATGGAATCAAAGCTAACTTCAAAATTCGCCACAA<br>CATTGAAGATGGATCCGTTCAACTAGCAGACCATTATCAACAAAATACTCC<br>AATTGGCGATGGCCCTGTCCTTTTACCAGACAACCATTACCTGTCGACACAA<br>TCTGCCCTTTCGAAAGATCCCAACGAAAAGCGTGACCACATGGTCCTTCTTG |

| | |
|---|---|
| | AGTTTGTAACTGCTGCTGGGATTACACATGGCATGGATGAGCTCTACAAAT<br>GA |

Supplementary Table 6. A codon optimized homolog for *Oxyrrhis marina* of the Rifampin ADP-ribosylating transferase from the bacterium *Citrobacter freundii*

| Name | Sequence |
|---|---|
| *arrO* | ATGGTGAAGGATTGGATCCCGATCTCTCACGATAACTACAAGCAGGTGCAG<br>GGACCGTTCTACCACGGAACCAAGGCGAACTTGGCGATCGGAGATTTGCTG<br>ACCACCGGCTTCATCTCCCACTTCGAGGACGGACGTATCCTGAAGCACATCT<br>ACTTCTCCGCGTTGATGGAGCCGGCTGTGTGGGGAGCTGAGCTGGCTATGTC<br>GCTGTCTGGCTTGGAGGGACGTGGCTACATCTACATCGTGGAGCCGACCGG<br>ACCGTTCGAGGACGATCCGAACCTGACCAACAAGAAGTTCCCGGGCAACCCG<br>ACCCAGTCCTACCGCACCTGCGAGCCGTTGCGCATCGTGGGCGTGGTGGAGG<br>ACTGGGAGGGACACCCGGTGGAGTTGATCCGTGGAATGTTGGACTCGTTGG<br>AGGACTTGAAGCGCCGTGGCTTGCACGTCATCGAGGATTAG |

Supplementary Table 7: A codon optimized homolog for *Oxyrrhis marina* of the Rifampin ADP-ribosylating transferase from the bacterium *Citrobacter freundii,* and the intergenic region between rhodopsin tandem repeats of *O. marina* at its 5'-end (which is shown in lowercase)

| Name | Sequence |
|---|---|
| *arrO-N* | aattttgggagttgggctggaagatggggttggtggggatcgggggagaggtgactggtgtgtggtcg<br>agATGGTGAAGGATTGGATCCCGATCTCTCACGATAACTACAAGCAGGTGCA<br>GGGACCGTTCTACCACGGAACCAAGGCGAACTTGGCGATCGGAGATTTGCT<br>GACCACCGGCTTCATCTCCCACTTCGAGGACGGACGTATCCTGAAGCACATC<br>TACTTCTCCGCGTTGATGGAGCCGGCTGTGTGGGGAGCTGAGCTGGCTATG<br>TCGCTGTCTGGCTTGGAGGGACGTGGCTACATCTACATCGTGGAGCCGACCG<br>GACCGTTCGAGGACGATCCGAACCTGACCAACAAGAAGTTCCCGGGCAACCC<br>GACCCAGTCCTACCGCACCTGCGAGCCGTTGCGCATCGTGGGCGTGGTGGAG<br>GACTGGGAGGGACACCCGGTGGAGTTGATCCGTGGAATGTTGGACTCGTTG<br>GAGGACTTGAAGCGCCGTGGCTTGCACGTCATCGAGGATTAG |
